# Non-canonical, ligand-independent basal mGluR1 signaling tunes Kv1.2 surface expression through PKA

**DOI:** 10.1101/2025.11.12.687507

**Authors:** Sharath C. Madasu, Anthony D. Morielli

## Abstract

Voltage gated potassium channels are major determinants of excitability of Purkinje cell (PC) neurons in the cerebellum. We have previously shown that in the cerebellum, activation of mGluR1 with agonist DHPG leads to reduced surface expression of Kv1.2 and that Kv1.2 co-immunoprecipitates with PKC-γ and CaMKII which are known interactors of mGluR1. However, mGluR1 can also signal independently of agonist through a constitutively active, protein kinase A-dependent pathway. Here we show that in HEK293 cells, co-expression of mGluR1 increases the surface expression levels of Kv1.2. This effect occurs in absence of mGluR1 agonists and in the presence of an mGluR1 inhibitor. Co-expression of known downstream effectors of the agonist driven mGluR1 pathway, including PKC-γ and CaMKII, had no effect on Kv1.2 surface expression or on the ability of mGluR1 agonist to modulate that expression. In contrast, the inverse agonist BAY 36-7620 significantly reduced the mGluR1 effect on Kv1.2 surface expression, as did pharmacological inhibition of PKA with KT5720.

## Introduction

The voltage gated potassium channel Kv1.2 plays an important role modulating membrane potential of neurons. A major role of Kv1.2 within the cerebellum is to modulate the excitatory (1) and inhibitory (2) input to Purkinje cells (PC). Excitatory input to PCs is influenced by Kv1.2 expressed PC dendrites, which receive excitatory input from parallel fibers. Khavandagar *et al* have shown that Kv1.2 prevents random Ca^2+^ spikes and dendritic hyper excitability of Purkinje cells(1), they also show that blocking Kv1.2 potentiated and prolonged the responses to parallel fiber stimulation. McKay *et al* showed that Kv1 channels shape the CF firing onto PC, inhibiting Kv1 channels resulted in spontaneous firing of CF suggesting Kv1 channels are expressed in presynaptic sites as well. On PC, blocking Kv1 channels reduced the current threshold to evoke Ca-Na burst and altered the firing pattern of Purkinje cells, from firing trains of action potentials, PC with Kv1 channels blocked fired brief and rapidly inactivating action potentials(3). Inhibitory input to PCs is influence by Kv1.2 expressed in basket cell axon terminals, which provide strong inhibitory input to Purkinje cells (2). PCs are the major information processing units of the cerebellum and provide the sole output of the cerebellar cortex, placing Kv1.2 in a central position to influence information processing in the cerebellum(4).

We have previously shown that, in the cerebellum, suppression of Kv1.2 with the specific inhibitor Tityus toxin (TsTX) or by secretin receptor mediated endocytosis of Kv1.2 results in enhanced EBC (2,5). We have also shown that EBC affects surface expression of Kv1.2 (5,6). mGluR1 is a G-protein coupled receptor which is essential for normal cerebellar function and for EBC activity(7). We have previously shown that in the cerebellum mGluR1 activation leads to loss of surface Kv1.2 (Chapter 2). The cellular mechanisms by which mGluR1 modulates Kv1.2 are not well understood. In a previous study we used mass spectrometry to show that that Kv1.2 physically interacts with potential regulatory proteins known to be activated by mGluR1, including PKC gamma, CaMKII, Grid2, and G_q/11_. We also found that mGluR1 regulates Kv1.2 in part through a PKC-dependent mechanism. In this study we used HEK293 cells and co-expression of mGluR1 and selected components of its downstream signaling pathways as a way to analyze the molecular mechanism by which mGluR1 regulates Kv1.2 in greater detail. Interestingly, we found that mGluR1 activation with the agonist DHPG had no effect on Kv1.2 surface expression in HEK cells. This lack of effect on Kv1.2 persisted even with co-expression of mGluR1 downstream effectors that are endogenously expressed in cerebellar neurons but not in HEK293 cells, including PKC-γ, Grid2 or CaMKIIα, either individually or in combination. In contrast, expression of mGluR1 elicited a strong increase in Kv1.2 surface expression. This phenomenon was completely independent of agonist activation of the receptor but was blocked using the inverse agonist BAY-36-7620. Indicating that mGluR1 is constitutively active.

In heterologous expression system it has been shown that mGlur1a and mGluR5 have basal agonist independent activity. mGluR1a is shown to increase both inositol phosphate (IP) and cAMP levels in HEK cells without activation. This constitutive activity was attenuated in N782I and E783Q mutants (8). It has also been shown that cAMP – PKA prevents desensitization of mGluR1a, 1b by preventing internalization by GRK2 (GPCR kinase 2) and arrestin 2 (9), therefore potentiating the agonist independent signaling of the receptor (10). Similarly Homer proteins are intracellular scaffolding proteins that interact with C-terminal PPXXF domains of receptors and act as a scaffold for other signaling proteins. Homer proteins are classified into long form which include homer 1b, 1c,2 and 3 proteins while Homer 1a is the short form. Homer 3 protein is known to interact with mGluR1/5 and prevents constitutive activation (11) while Homer1a binds competitively with mGluR1 replacing Homer3 and activates the receptors. In the cerebellum mGluR1 receptors were shown to be constitutively active via PKA and increase the intrinsic excitability of PCs by affection HCN ion channels (12).

Here we show that mGluR1 is constitutively active and signals via PKA to increase surface Kv1.2 because PKA inhibitor KT-5720 prevented mGluR1 mediated increase in surface Kv1.2

## Methods

### Materials

Flag-mGlur1α plasmid are a gift from Dr. Stephen Ferguson, Canada, CaMKII GFP was a gift from Dr. Stephen Tavalin, University of Tennessee, USA. Grid2 plasmid was a gift from Dr. Carol Levenes, Centre de Neurophysique, Physiologie et Pathologie, France.

### Cell culture and transfection

HEK – M cell line which stably expresses Kv1.2/Kvβ2 was used for all the experiments. Cells were maintained in DMEM-F12 medium supplemented with 10% FBS, Pen/Strep glutamine, Zeocin, G418 and were maintained in 5% CO_2_ environment at 37°C. On the day of transfection 35 mm dishes were coated with 1µg/ml PEI for 15 min followed by wash with PBS. The HEK cells were plated at a medium confluence on to the PEI coated dishes and were let grow to achieve 70-80% confluence. Medium was replaced on the day before transfection.

Required amount of DNA (maximum of 5µg) was mixed with 150mM NaCl incubated by 10 incubation. 18µl (1µg/ml) of PEI was mixed with 112µl of 150mM NaCl and was incubated for 10 min followed by mixing with the plasmid mixture and another 10 min incubation. Cells were then transfected with the PEI-plasmid mixture and were allowed to grow for 24hours.

### Flow cytometry

Transfected cells were plated on to a PDL coated (1mg/ml) 6 well dish and were serum starved in Neurobasal-A medium overnight. The next day medium was replaced at least 1hour before drug treatment. All drugs were added from stock solution made with Neurobasal medium and were added for indicated times. At the end of drug incubation endocytosis was stopped by addition of 0.4% sodium azide and cells were incubated for 20 min at 37°C in the incubator. Cells were then scrapped, washed with 0.1% FBS in PBS solution (flow sauce). The cells were resuspended and incubated in anti-Kv1.2 extracellular antibody as described in (13) for 1 hour. The cells were then washed once and anti-rabbit SPRD antibody was added and incubated for 17 min. The cells were then washed twice in flow sauce spun down and resuspended in 300µl of flow sauce with 2µg of DAPI for live/dead cell staining. The stained cells were then enumerated on MACS VYB benchtop flow cytometer (Milteny biotech). Initially 10,000 events were counted, as the number of transfections increased 100,000 events were counted as indicated.

### Western Blots

Cells were transfected, and serum starved as above and were lysed in RIPA buffer with the following formulation: 50mM Tris, 150mM NaCl, 2mM EDTA, 0.25% deoxycholate, 1% NP40 alternative, 10%glycerol, 1mM DTT, supplemented with 1x HALT protease inhibitor, 100mM Na_3_VO_4_, 1mM NaF, 5mM BAPTA, BPvphen, mg132. The lysates were subjected to centrifugation to separate nuclear pellet and were separated on SDS – PAGE and was transferred on to nitrocellulose paper and subjected to western blot analysis. Total Kv1.2 was detected via anti-Kv1.2 antibody from Neuromab, anti-mGluR1, anti PKA ser/Thr substrate antibodies from CST.

### Immunofluorescence microscopy

Immunofluorescence imaging was performed as described previously(14) using anti flag anti body (sigma). Images were acquired and processed with the DeltaVision deconvolution restoration microscopy systems (applied Precision, Issaquah, WA).

## Results

### Expression of functional mGluR1 receptors in HEK293 cells

HEK-M cells were transfected with (Flag-mGluR1), encoding mGluR1. Expression of mGluR1 protein was verified by immunoblot and immunofluorescence. Western blot analysis revealed expression of FLAG-mGluR1 protein was stable between 24 to 72 hours post-transfection (Figure 1A). Immunofluorescence analysis showed no detectable mGluR1 in un-transfected cells, i.e. those not expressing GFP (Figure 1B). mGluR1 expression in transfected cells was detectable at the cell surface by using an antibody directed at the FLAG-tag in the extracellular N-terminus of the expressed mGluR1 (Figure 1C).

**Figure 1:**
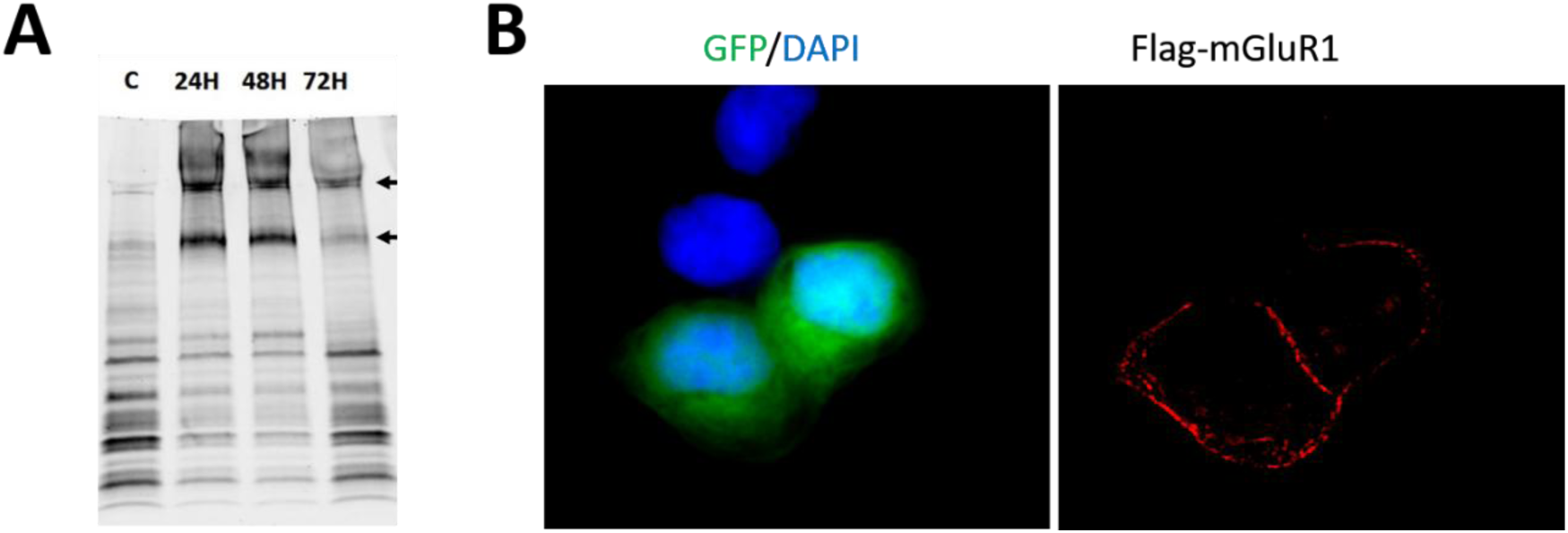
Expression of mGluR1 in HEK cells. A) Western blot of cell lysates transfected with control vector (PRK5) in lanes C, mGluR1 and lysates collected 24 hours (24H), 48 hours (48H), 72 hours (72H) post transfection. Arrows indicate the monomer (around 150 Kd and dimers 250 Kd). B Immunofluorescence confocal microscopy images of HEK cells co-transfected with Flag-mGluR1 and GFP, counterstained with DAPI nuclear stain, only cells transfected with GFP (green) showed surface mGluR1(red) expression.

### mGluR1 agonist DHPG activates calcium signaling and increases ERK phosphorylation

We next asked if the expressed receptors were functional. Augustine *et al* have shown that mGluR1 increases ERK phosphorylation (15). Several researcher have shown that mGluR1 activation leads to release of intracellular calcium (16,17). In HEK-M cells expressing mGluR1, application of DHPG increased ERK phosphorylation only in cells transfected with mGluR1 (Figure 2A), while in cells pre-incubated with mGluR1 noncompetitive antagonist YM 298198, the inhibitor blocked the increase in phosphorylation). Similarly, DHPG (100µM) and glutamate (200µM) caused a transient increase in intracellular calcium measured in individual cells using the calcium indicator dye Fluo4-AM (Figure 2B). Preincubation with the competitive antagonist YM 298198 completely blocked the calcium signaling events in HEK cells transfected with mGluR1 (Figure 2B) YM 298198 did not affect calcium signal detected by application of the calcium ionophore A23187. We also observed that neither glutamate nor DHPG were able to alter intracellular calcium after an initial application, consistent with desensitization of the mGluR1 receptor response(9,18). Collectively, these findings confirm the expression of functional, agonist sensitive mGluR1 receptors in HEK-M cells.

**Figure 2:**
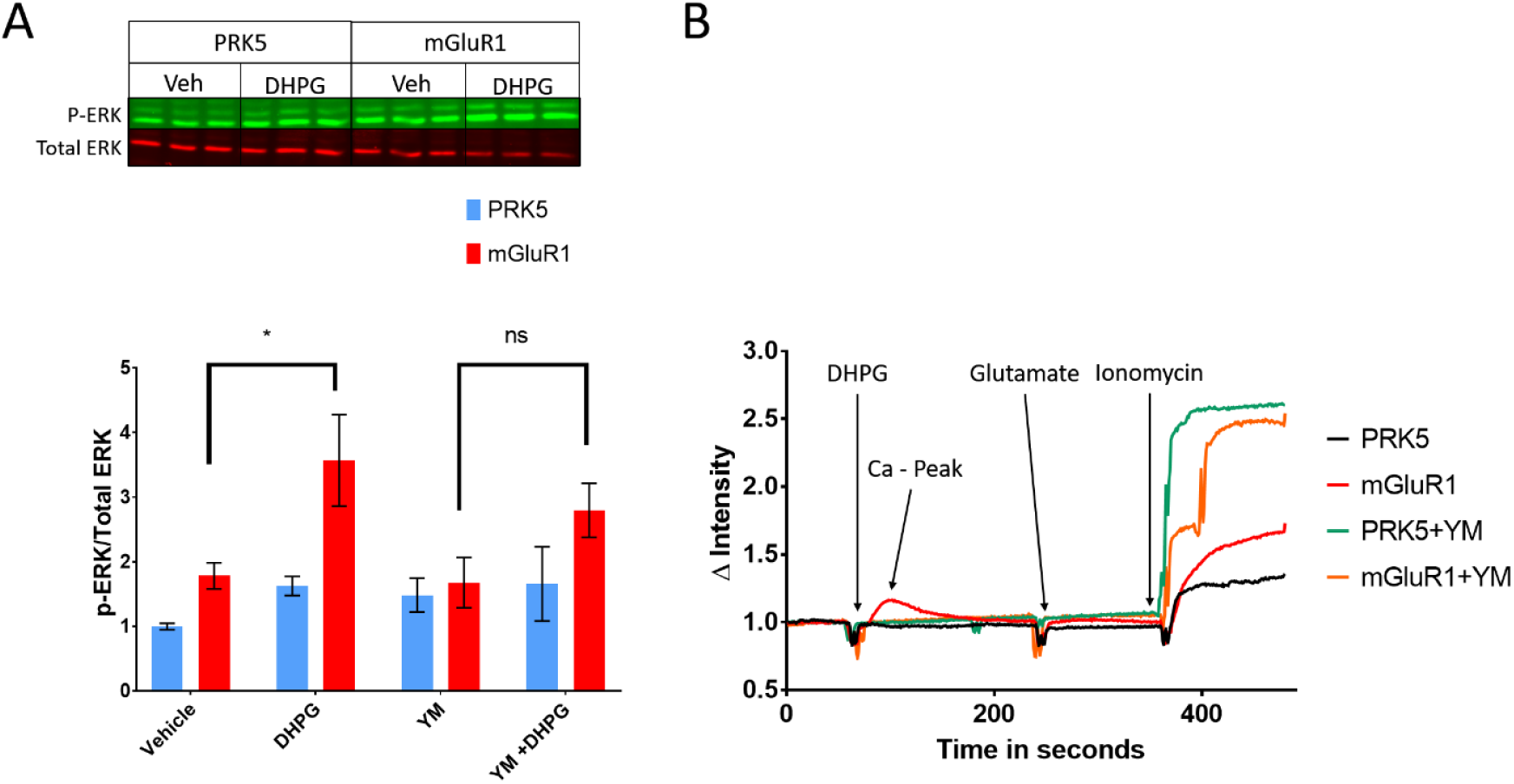
Functional expression of mGluR1 in HEK cells. A) Western blot of cell lysates transfected with control vector (PRK5), mGluR1. Cells were in glutamate free medium and were incubated in HBSS for 1 hour followed by treatment with DHPG (100µM) for 10 min. Cell were incubated with YM 298198 (YM in figure) prior to DHPG. The lysates were then collected. B) HEK cells co-transfected with Flag-mGluR1 and were in glutamate free overnight and were incubated with Flou-4AM Cells transfected as indicated were treated with DHPG or 200 µM glutamate. Only cells transfected with mGluR1 and treated with DHPG or glutamate (DHPG trace is highlighted) show increase in calcium levels (Ca-Peak red line). mGluR1 inhibitor YM 298198 (YM in figure legend) blocks Calcium increase.

### Detection of Surface Kv1.2 by Flow Cytometry

In our previous studies we have shown that Kv1.2 is regulated by trafficking of Kv1.2 from cell surface via endocytosis and that such endocytosis results in reduced Kv1.2 ionic current. In previous studies flow cytometry was used to study the surface expression of Kv1.2 in HEK cells(13,14). Those studies have identified changes in surface expression in response to activation of another G-protein coupled receptor, the M1 muscarinic receptor (19,20). Therefore, in this study we also used flow cytometry to detect mGluR1 effects on Kv1.2 surface trafficking. To accomplish this, HEK cells were co-transfected with Flag-mGluR1, eGFP, and control vector PRK5 plasmids. Live cells were distinguished by exclusion of DAPI nuclear stain, and GFP expression was used as positive marker for transfection. Only cells that were live and had GFP expression were selected for analysis of surface Kv1.2. Surface Kv1.2 was detected using a fluorescently labeled antibody directed against an extracellular epitope within Kv1.2 ((13), Methods). Cells that received PRK5 only, received 2^0^ Ab only and i.e. do not express Kv1.2, were negative controls for antibody staining (Figure 3).

**Figure 3.**
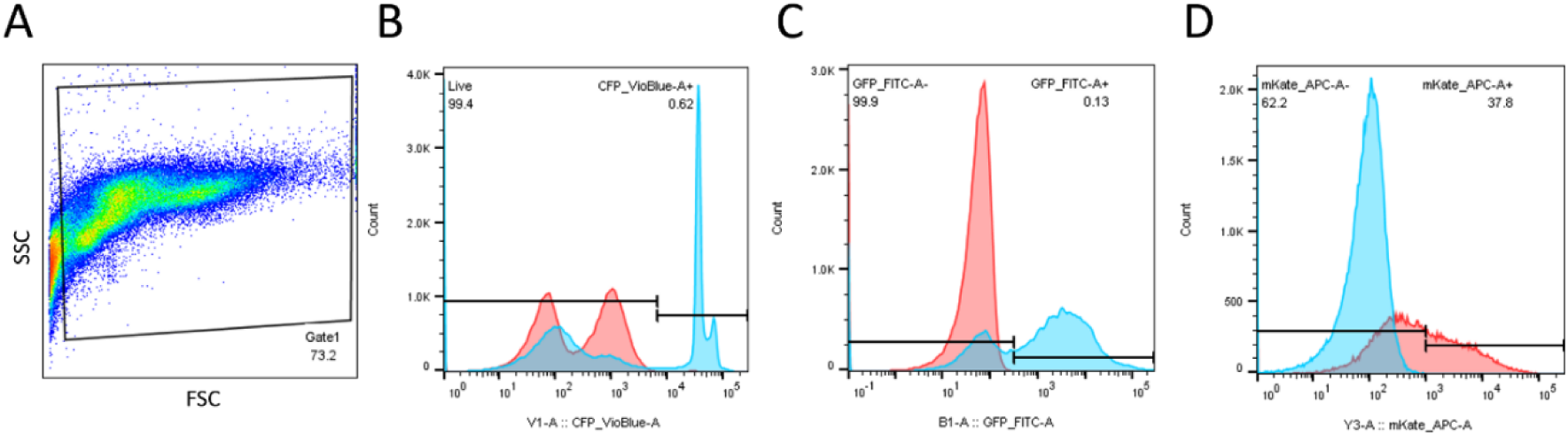
Gating strategy for estimating surface Kv1.2 via flow cytometry. A) Side scatter (SSC) and forward scatter (FSC) profile of HEK cells. B) Live cell gating. DAPI was used as Live dead marker, PRK5 only transfected cells are DAPI-ve (red shaded population) was used as negative control. mGluR1 transfected cells (blue shaded population). C) PRK5 only cells (red shaded population) was used to select transfected cells which are GFP +ve (blue shaded population.). D PRK5 (blue shaded) only cells which are -ve for Kv1.2 antibody was used to gate Kv1.2 +ve cells (red shaded cells).

**Figure 4.**
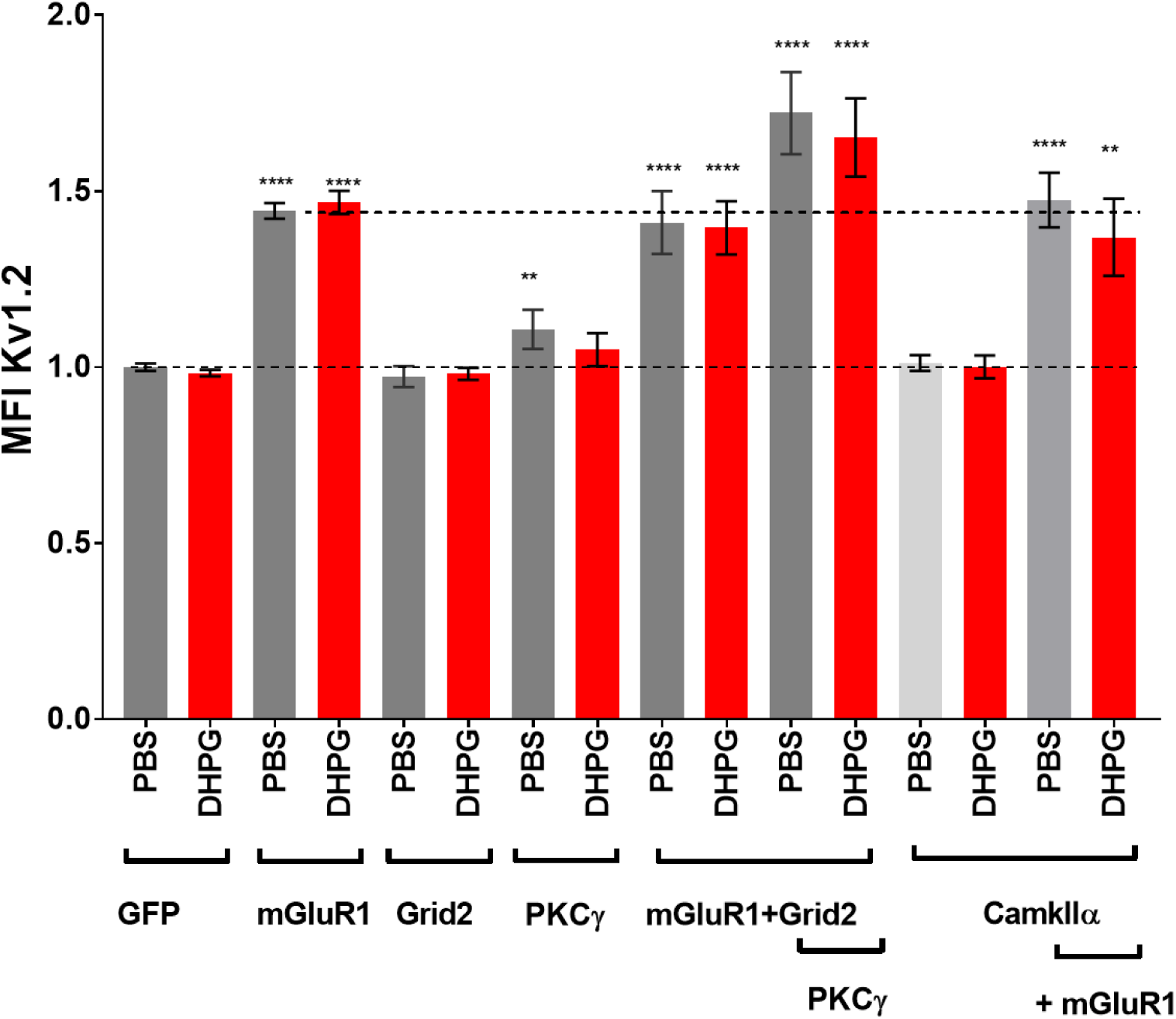
mGluR1 increases surface expression of Kv1.2: Hek 293 M1 cells that stably express Kv1.2 and β2 sub units were transfected as mGluR1, PKCγ, Grid2, CaMKIIα (GFP tagged) alone or in combinations indicated in figure. The cells were analyzed via flow. Median fluorescence intensity was normalized to GFP transfected PBS treated cells. **** indicates P<0.0001 vs GFP PBS cells, ** indicates p=0.0031 vs GFP PBS cells. P value calculated via Tukey’s multiple comparison test 2-way Annova with transfections and Drug treatments as variants. CaMKIIα were compared via one-way annova and Sidak’s multiple comparison test as the transfections were performed separately.

### mGluR1 activation by agonist does not affect Kv1.2 surface expression

In a previous study we found that mGluR1 activation decreases the net expression of Kv1.2 at the cell surface in cerebellar slices. We therefore expected mGluR1 activation in HEK-M cells to have a similar effect on Kv1.2 surface levels. As shown in Figure 5A, the mGluR1 agonist DHPG did not reduce the surface expression of Kv1.2 as we observed in the cerebellum. This lack of effect was likely not caused by desensitization of mGluR1 for several reasons. First, as shown in Figure 2, both MAP kinase and calcium signals were observed in response to DHPG treatment in HEK cells expressing mGluR1. Second, DHPG was also without effect in cells that had been pre-treated with Sodium pyruvate and glutamate pyruvate transaminase (GPT) to scavenge of any glutamate that might have accumulated in the bath from the cells themselves (Figure 5b).

**Figure 5.**
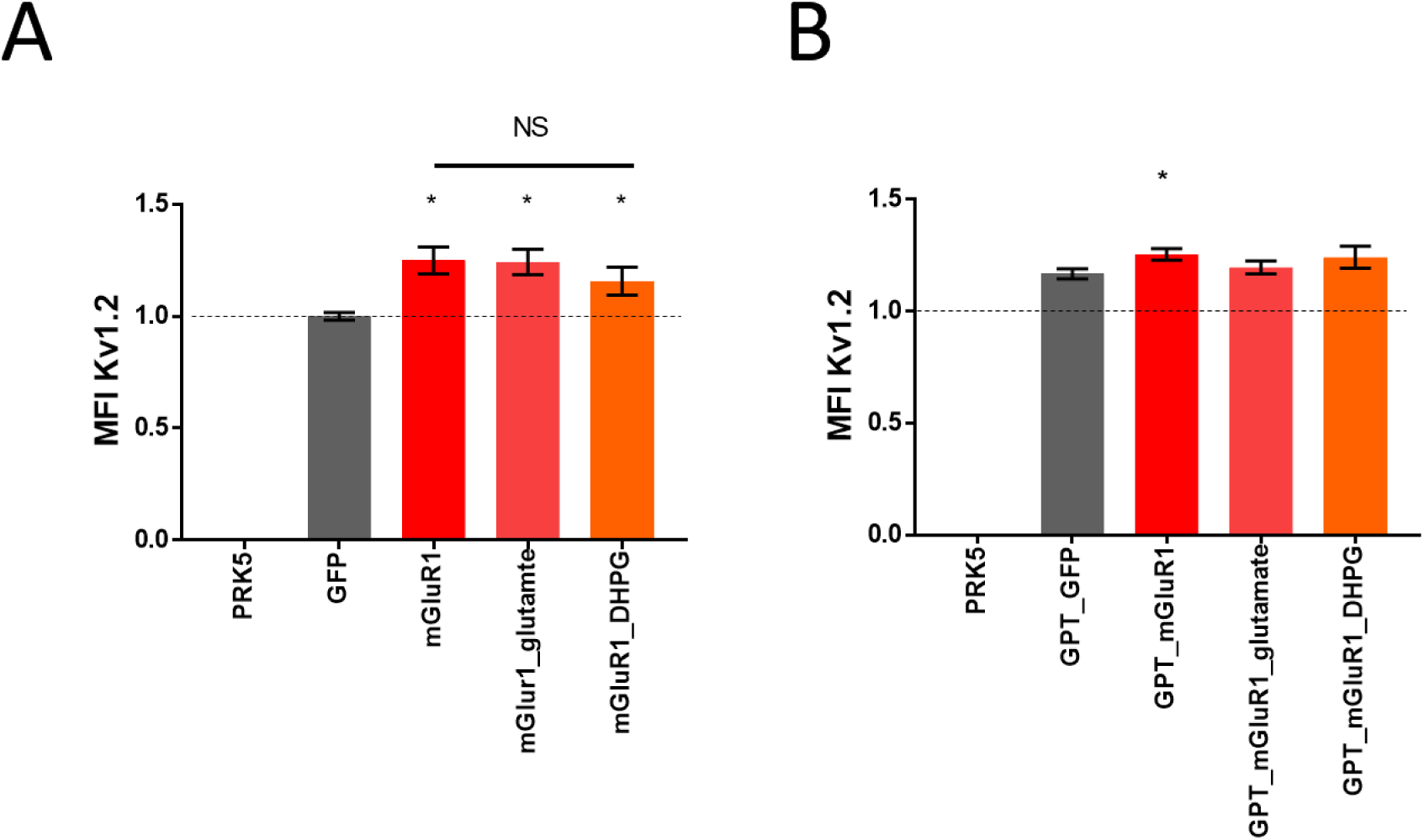
mGluR1 increases surface expression of Kv1.2 independent of Agonist: Hek 293 M1 cells that stably express Kv1.2 and β2 sub units were transfected as mGluR1. The transfected cells were then serum starved in neurobasal medium without glutamate (A) and Cells in (B) were treated with glutamate pyruvate transaminase and 100mM sodium pyruvate (GPT in B) for 2 hours followed by treated with 100µM DHPG or Glutamate for 10 min. MFI was analyzed via flowcytometry and normalized to GFP transfected cells. PRK5 cells are mock transfected and are Kv1.2 Ab (-) ve, GFP (-) ve and were used to gate transfected cells. * indicates p<0.05 vs GFP, p value calculated by students T test with Welch correction.

### Over expression of PKC-γ, Grid2, CaMKIIα does not affect surface expression of Kv1.2

HEK cells do not express many of the proteins known to be involved in mGluR1 signaling in the cerebellum. We have identified mGluR1 interacting proteins that bind to Kv1.2 such as PKC γ, CaMKIIα, and Grid2. None of these proteins are expressed endogenously in HEK cells. We therefore transfected Kv1.2/β2 stable M1 cell lines with PKC-γ, Grid2 or CAMKII, alone or in combination, in an attempt to reconstitute the mGluR1 signaling complex in HEK cells. We did not observe any change in level of surface expression of Kv1.2 in the cells transfected with Grid2 and CaMKII compared to control transfected cells Figure 4, while cells transfected with PKC-γ had a slight increase in surface Kv1.2 (p=0.0031 vs GFP PBS cells). In no instance did expression of these proteins, alone or in combination, result in Kv1.2 modulation by DHPG (Figure 4).

### Modulation of Kv1.2 via constitutively active mGluR1 or basal mGluR1 activity

Although DHPG had no effect on Kv1.2 surface expression despite the presence of functional, agonist sensitive mGluR1 receptors, we consistently observed that mGluR1 expression alone elicited a significant increase in surface Kv1.2 levels relative to HEK cells expressing Kv1.2 but not mGluR1. Like many G-protein coupled receptors, mGluR1 is capable of signaling in the absence of agonist activation (8). We therefore hypothesized that such constitutive activation of mGluR1 or basal activity was responsible for the elevation in surface Kv1.2 observed in cells co-expressing mGluR1. To test this idea, we used an inverse agonist, BAY 36-7620, which blocks constitutive activity of mGluR1(21,22). We found that incubation with Bay 36-7620 overnight significantly attenuated the mGluR1 mediated decrease in surface Kv1.2 (Figure 6A).

**Figure 6.**
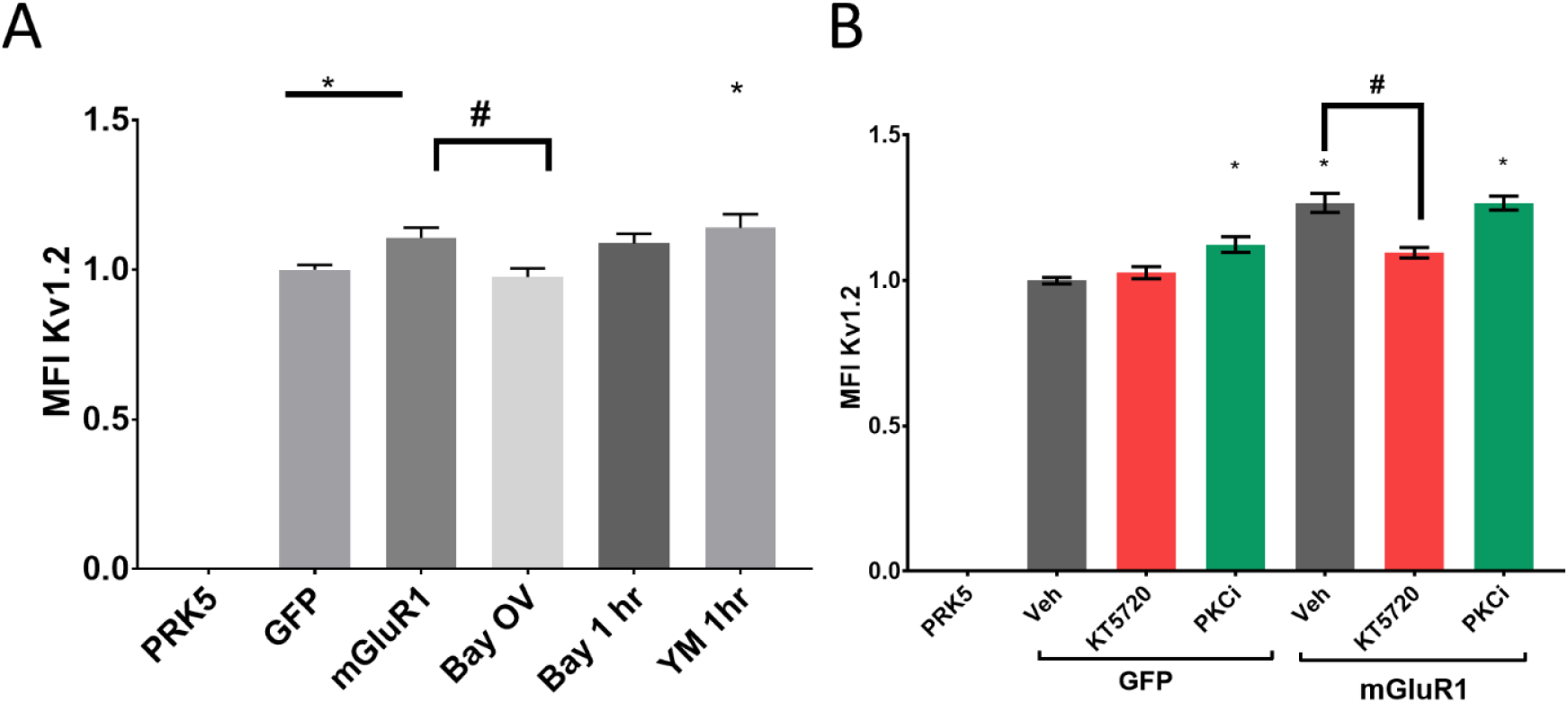
Constitutively active mGluR1 increases surface expression of Kv1.2 via PKA: Hek 293 M1 cells that stably express Kv1.2 and β2 sub units were transfected with mGluR1. The transfected cells were then serum starved in neurobasal medium without glutamate. A) Cells were incubated with mGluR1 inverse agonist BAY 36-7620 (BAY) for 1 hour or overnight (OV), with noncompetitive antagonist 100µM YM298198 for 1 hour prior to flow cytometry B) Cells were treated with PKC inhibition Go-6987 (1µM) and PKA inhibitor KT-5720 (5µM) 1 hour prior to flow analysis. * indicates p<0.05 vs GFP, p value calculated by students T test with Welch correction # indicates p<0.05 compared to mGluR1 Veh.

### mGluR1 increases surface Kv1.2 via PKA mediated mechanism and not via PKC

Several lines of research showed that mGluR1 also couples to and signals via G_s_ – PKA pathway in heterologous expression systems and in the cerebellum (8,12,23), intracellular loops 2 and 3 play an important role in binding of mGluR1 to Gs G – proteins (8). The same study also showed that mGluR1 transfection results in an increase in basal levels of cAMP and inositol phosphate production(8). Further it has been shown that PKA activation prevents mGluR1 receptors from regular constitutive internalization therefore potentiating the constitutive activity and signaling inositol pathways (9). In the cerebellum it was shown that mGluR1 via PKA increases intrinsic excitability by modulating HCN potassium channels(12). We hypothesized that mGluR1 regulates Kv1.2 in HEK cells via PKA when constitutively active. To test this hyothesis, we incubated the cells with PKA inhibitor KT 5720 and PKC inhibitor Go6980 for 1 hour prior to flowcytometry. In cells expressing Kv1.2 but not mGluR1, KT 5720 did not affect surface Kv1.2, while PKC inhibitor increased surface expression of Kv1.2 (1.124±0.085 vs GFP veh, p=0.0043). However, in cells co-expressing mGluR1 along with Kv1.2, KT5720 decreased the surface expression (1.26±0.13 mGluR1 vs 1.096±0.071 p<0.0001,) (Figure 6B). Collectively, these findings indicate that in HEK293 cells, constitutive activation of PKA signaling by mGluR1 elevates surface Kv1.2 levels.

## Discussion

Here we identify a new mechanism for the regulation of Kv1.2 homomeric ion channels in a heterologous expression system. We have also shown that in the cerebellum Kv1.2 interacts with several proteins including PKC-γ, CaMKIIα, G_q/11_ etc. which are also known to interact with Kv1.2(24). In that study, agonist activation of mGluR1 reduced the level of Kv1.2 at the cell surface in cerebellar slices . In depth analysis of the molecular mechanisms of that modulation in native tissue is complicated by the presence of an array of active signaling systems, each potentially able to modulate Kv1.2. This natural complexity also makes it difficult to determine whether the effect of mGluR1 on Kv1.2 is direct or is perhaps secondary to mGluR1 effects on availability of other neurotransmitters capable of modulating the channel. Numerous studies use HEK293 cells as an experimental system for studying Kv1.2 regulation in isolation of such confounding factors. We therefore in this study we attempted to reconstitute in HEK293 cells the mGluR1 agonist-driven modulation of Kv1.2 that we observed in cerebellar tissue.

In HEK cells, mGluR1 activation with the specific agonist DHPG or with natural agonist glutamate, did not affect surface Kv1.2 levels. This is in contrast to the effect we observed in cerebellar slices where DHPG application reduced surface Kv1.2. This lack of effect on surface Kv1.2 in HEK cells was not because the mGluR1 receptors were non-functional since DHPG did elicit an increase in ERK phosphorylation and could elevate intracellular calcium (Figure 2A).

It is possible that although mGluR1 can couple to some signaling pathways in HEK cells, the pathways linking the receptor to Kv1.2 might be missing. In our previous work we identified several proteins that interact with Kv1.2 in the cerebellum but that are not endogenously expressed in HEK cells. These include Grid2 (24,25) an ionotropic glutamate ion channel which does not respond to glutamate directly but does so when mGluR1 is activated via DHPG or glutamate, as well as (22,23) PKC-γ and CaMKIIα, also known interactors of mGluR1 (24,26). We anticipated that co-expression of these proteins might reconstitute an ability of mGluR1 to regulate Kv1.2 in an agonist-dependent way. As shown in Figure 4, this did not occur. However, although Grid2, PKC-γ, CaMKIIα are good candidates for reconstituting mGluR1 signaling to Kv1.2 in HEK cells, they are not the only proteins that interact with cerebellar Kv1.2, proteins such as DLG (Disk Large Homologs), 14-3-3, WNK/STK39, Spectrin class proteins, adapter protein 2 and many others were shown to interact with Kv1.2. It is therefore possible that co-expression of other proteins, in particular scaffold proteins such as Disk large proteins (also known as post synaptic density, PSD) interact with both mGluR1 and Kv1.2 or isoforms of Homer proteins that are known to regulate constitutive activation, might lead to successful reconstitution mGluR1 signaling to Kv1.2.

Although mGluR1 agonist-dependent signaling did not affect Kv1.2, we were surprised to find that expression of mGluR1 alone was able to affect surface Kv1.2 levels. Agonist-dependent mGluR1 signaling proceeds, in part, through Gq/11 (27). However, like many G-protein coupled receptors, mGluR1 can signal in the absence of agonist. Such constitutive activation or basal activity is, in the case of mGluR1, could be mediated by coupling of the receptor to Gs and a resultant increase basal levels of cAMP and PKA (8,10,11,23). Connors *et al* (13) had previously shown that both cAMP and PKA can modulate surface Kv1.2 levels in HEK cells. Hence to determine if mGluR1 increases surface expression via a cAMP/PKA-dependent pathway we inhibited both PKA and PKC. In our current study only PKA inhibitor KT5720 reduced the mGluR1 mediated increase in surface Kv1.2, suggesting that PKA is largely responsible for the increase. It is important to note that the effect of PKA on Kv1.2 is not straightforward. In the study by Connors *et al*, indirect evidence suggested that PKA could, depending on its level of activity, reduce surface Kv1.2 levels. However constitutive mGluR1 signaling involves not only PKA but involve other components, including Homer proteins, arrestins and phospholipase C(28). Thus, although the effect of mGluR1 on Kv1.2 appears to require PKA activity, the specific outcome of an increase in surface Kv1.2 levels likely involves these other signaling pathways as well.

Although mGluR1 constitutive activity has been shown in the cerebellum, whether or it affects Kv1.2 there remains to be determined. In this respect it is important to note that in this study we examined Kv1.2 homomultimeric channels. However, in the cerebellum Kv1.2 is thought to exist as homomultimers, but also as heteromultimers with Kv1.1 and possibly other Kv1 alpha subunits (9,29). These subunits could confer different sensitivities to mGluR1 constitutive activity. Conversely, they might be the missing component required to couple Kv1.2 to agonist-dependent mGluR1 signaling. For example, the Kv1.1 subunit is dephosphorylated when mGluR1 receptors are activated (30) and the extent of inactivation of the Kv1.1/β1.1 ion channels was reduced in HEK cells. Direct activation of PKA phosphorylates intracellular Kv1.1 and promotes the surface Kv1.1 levels, while direct activation of PKC stimulated protein synthesis of Kv1.1 in HEK cells (31) and in cerebellar granule cells, inhibition of PKC by bisindolylmaleimide reduced the expression of Kv1.1 (32) but there was no evidence of direct phosphorylation (31). In contrast Kv1.3 and Kv1.4 are downregulated by PKC activation (33,34). Therefore, while the current study provides enticing new insight into the mechanisms by which mGluR1 might regulate Kv1.2, the actual mechanisms for regulating the channel are likely to be strongly influenced by the signaling systems at play in specific neurons.

